# Exposure-driven modulation of neural response to novel word forms in toddlers

**DOI:** 10.64898/2026.11.25.769138

**Authors:** Pia Rämä, Oytun Aygün

## Abstract

An enhancement of an early brain signal in response to novel word forms has been associated with lexical learning in adults. Here, we measured neural responses to novel pseudowords in 24-month-olds to investigate whether a similar early modulation occurs in young children. Participants were presented with familiar French words (e.g., *bateau*, “boat”) and novel pseudowords (e.g., *bamon*). Event-related potentials were time-locked to the word disambiguation point (i.e., the onset of the second syllable) and analyzed across three time bins (∼ 4 minutes each) during the experiment. We observed an early positive response, P100, occurring around 100 ms after the word disambiguation point, which decreased for pseudowords over the course of repeated exposure. No such amplitude change was observed for familiar words. These results demonstrate an early reduction in neural responses to novel word forms, suggesting rapid and dynamic changes in brain activity associated with the familiarization of novel lexical items.

## Introduction

Early vocabulary development is marked by rapid changes in both behavioral and neural processes as infants begin to acquire and produce new words. Vocabulary growth, in particular, accelerates during the second year of life, and around two years of age toddlers typically acquire new words at a striking pace. Event-related potential (ERP) studies show that both age and vocabulary size modulate neural responses to familiar and unfamiliar words in infancy (e.g., Mills et al., 1993, 1997; Aygün & Rämä, 2022). Word familiarity effects are typically observed in a time window of approximately 200–500 ms following word onset while an earlier P100 component appears to reflect age-related latency differences that are independent of word familiarity (i.e., known versus unknown words, cf. Mills et al., 2004). The N200–500 response in younger infants (13–17 months) is more broadly distributed across the scalp than in older infants (around 20 months) (Mills et al., 1993, 1997), suggesting increasing functional specialization of word-related neural activity with development. Consistent with this interpretation, ERPs elicited by consonant mispronunciations differ from those elicited by correctly pronounced known words in 20-month-olds, whereas 14-month-olds do not yet show a reliable distinction between these word forms (Mills et al., 2004).

Previous studies examined neural representations of already known words and there are only a small number of studies that have shown that ERP responses can change as a function of newly learned word meanings acquired during the experiment (Mills et al., 2005; Torkildsen et al., 2008). For instance, the amplitude of the N200– 500 component in response to words paired with specific objects increased following training (Mills et al., 2005). In another study, infants were first trained on novel word– object associations. During the test phase, these newly learned associations were then violated by pairing the objects with novel words. This violation elicited an N400 effect, characterized by greater negativity for incongruent than for congruent pairs, particularly in infants with larger vocabularies (Torkildsen et al., 2008). Together, these findings suggest that the N200–500 and N400 components are sensitive to newly acquired word meanings and dynamically change as a function of learning during the course of an experiment in the infant brain.

Lexical information is processed much earlier in time in the adult brain (e.g., MacGregor, Pulvermüller, Van Casteren, & Shtyrov, 2012; Shtyrov & MacGregor, 2016). When adult participants are presented with written words (e.g., *‘lake”*) and meaningless pseudowords (e.g., *“lape”*), word-specific neuromagnetic responses can be detected as early as approximately 70 ms following word onset. A similar early enhancement has been observed 50–80 ms after the spoken word identification point, suggesting that lexical processing is extremely rapid also in the auditory modality (MacGregor et al., 2012). Moreover, even brief auditory exposure to meaningless pseudowords has been shown to induce changes in early neural responses (e.g., Kimppa et al., 2015, 2016; Leminen, Partanen, & Shtyrov, 2023). More specifically, when adult participants were presented with known Finnish words (e.g., *keto*, “meadow”) and pseudowords (e.g., *teto*), the neural response to pseudowords became larger than that to known words after approximately 30 minutes of exposure (Kimppa et al., 2015). Notably, the magnitude of this enhancement was associated with behavioral outcomes: better performance in spontaneous free recall and recognition tasks correlated with greater pseudoword response amplitudes, suggesting its role in word learning mechanisms.

A similar enhancement has also been observed in school-aged children, but it emerges much more rapidly—within approximately 4 minutes—compared to around 20–30 minutes in adults (Partanen et al., 2017). While response latencies for native-like stimuli were comparable in children and adults (approximately 50 ms), responses to non-native or non-speech stimuli (around 90 ms after stimulus onset) were observed only in children. Taken together, these findings suggest that early neural enhancement may reflect the formation of novel word representations, potentially indexing a cortical mechanism for rapid and automatic word learning. In adults, this mechanism appears to be phonology-dependent and supports the formation of neural memory circuits for newly learned word forms.

As noted above, by the end of the second year of life, toddlers undergo a rapid expansion of vocabulary skills, and explicit word learning in early childhood is typically associated with enhanced neural responses occurring approximately 200–500 ms after word onset. However, word learning in natural environments rarely involves explicit training: infants acquire words through everyday interactions with caregivers, including exposure to overheard speech (e.g., Akhtar, Jipson, & Callanan, 2001). In the present study, we examined whether 24-month-old toddlers exhibit early neural responses that are modulated by repeated exposure to meaningless words in the absence of explicit training. During the experiment, participants were presented with four novel pseudowords (e.g., *bamon*) and four familiar words (e.g., *bateau*, “boat”) in a passive listening task. ERPs were time-locked to the word disambiguation point (i.e., the onset of the second syllable), and amplitudes were analyzed across three consecutive time bins (each lasting four minutes) over the course of the experiment.

Based on toddlers’ word learning abilities and prior findings in adults and school-aged children, we hypothesized that neural response enhancement would emerge for pseudowords over time. Furthermore, if the magnitude of this enhancement reflects word-mapping abilities, and is thus related to vocabulary size, toddlers with larger vocabularies should exhibit greater enhancement than those with smaller vocabularies. In addition to early responses, we also examined the N200–500 component time-locked to the word recognition point, which has previously been associated with word familiarity and lexical-semantic processing in infants (Mills et al., 1993, 1997, 2005). We expected familiar words to elicit more negative responses than pseudowords across the experiment. Given the absence of explicit training on pseudoword meanings, we did not expect this later component to show modulation over time.

## Material and methods

### Participants

The final participant sample included twenty-three (12 girls, 13 boys) 24-month-old toddlers (range between 23 to 25 months, mean age 24 months). All participants came from monolingual French-speaking families in the Parisian region. They were recruited through a database of parents who previously volunteered to participate in child development studies. All children were born full term and no parents reported hearing or language impairment. Parents gave written informed consent before participation. Six other participants were tested but their data was rejected due to noisy electroencephalogram (EEG) recording (n = 5) or refusal to put on the EEG cap (n = 1). The French translation of the MacArthur-Bates Communicative Development Inventory for Words and Sentences (MCDI; Fenson et al., 1993) was used to measure productive vocabulary size. The parents were asked to fill the inventory at home during the two weeks following the study. The CDI was completed for 21 of 25 children in our final sample. The mean number of comprehended and produced words were 493 (SD = 135, range from 284 to 740) and 271 (SD = 102, range from 105 to 535), respectively. The study was conducted in conformity with the declaration of Helsinki and approved by the Ethics Committee of the University of Paris Descartes.

### Stimuli and the procedure

The audiostimuli consisted of two types of CVCV words: (a) French words (bateau (boat), canard (duck), chaussure (shoe), gateau (cookie)) and (b) pseudowords (bamon, cagnure, chautarre, gapeau). There were four tokens for each word. Mean durations of words and pseudowords were 328 ms and 324 ms, respectively. The durations of first and second syllables for words varied between 31 ms and 170 ms (mean 83 ms, SD 16) and 145 ms and 324 ms (mean 245 ms, SD 21), respectively. The durations of first and second syllables for pseudowords varied between 34 ms and 146 ms (mean 85 ms, SD 13) and 135 ms and 346 ms (mean 239 ms, SD 19), respectively. All words were recorded with a TASCAM Linear PCM Recorder (DR-07MKII). The speaker was a female native French speaker, and she pronounced the words slow and clear. The recordings were edited and normalized at 22 kHz sampling rate with Praat (5.3.19).

Before the testing, parents received a picture book that explained in a child-directed manner what happens during their laboratory visit (e.g., EEG capping, testing session). This book was meant to help both parents and children to be prepared for the testing session. After arriving at the laboratory, the children were allowed to play in the laboratory environment, and they were also familiarized with the EEG cap. During the experiment the child sat on the lap of a parent in a dimly lit room at 70 cm from a computer screen and two loudspeakers from which the sound was delivered. Parents were instructed not to communicate with the child during the experiment. Children were allowed to watch a silent cartoon on the screen throughout the experiment or to play with small toys positioned on the table in front of them during the experiment. Each word (4 familiar words and 4 pseudowords) was presented 20 times during the experiment. The presentation order was pseudo-randomised to ensure an even distribution of each word type during the experiment. The same word was never repeated more than twice. The mean stimulus onset asynchrony (SOA) was 1000 ms with a jitter ranging from 950 to 1050 ms. The experiment lasted about 12 minutes.

### EEG recording and pre-processing

Continuous electroencephalogram (EEG) was recorded (band-pass was 0.1 – 100 Hz, sampling rate was 250 Hz) from 128 electrodes using a Geodesic Sensor Net (GSN, NetStation EGIS V2.0). Impedances were kept below 50 kΩ. The data was filtered off-line (0.3–30 Hz) and segmented (700 ms, starting 100 ms before word recognition point). The epochs including artifacts (eye-movements, blinks, or motion artifacts) exceeding ±180 µV were excluded. The epochs with more than 30% of the channels contaminated were also rejected. Eye movement artifacts were removed with ocular artifact removal (OAR) tool (Gratton et al., 1983). Individual bad channels were replaced using spherical spline interpolation.

The data was divided into three equal experimental bins (early, middle, and late). Each bin consisted of 20 repetitions of each word, thus 80 repetitions of each word type, and lasted about four minutes. To obtain ERPs, the epochs were averaged separately for each subject, Word Type (word, pseudoword), and Phase (early, middle, and late). The averaged waveforms were re-referenced to the average of mastoids and baseline corrected. The averaged waveforms were grand-averaged across all participants. The mean number of accepted EEG epochs was 46, 44 and 41 for early, middle, and late phases for pseudowords, respectively, and 48, 44 and 41 for early, middle, and late phases for word, respectively. The mean number of accepted trials was not significantly different among the bins (One-way ANOVA: F(5, 149) = 0.56, p = 0.733).

### Data and statistics analyses

We calculated the amplitudes over two fixed time-windows after the word recognition point (50-150 ms, and 200-500 ms). The early window was chosen based on previous literature on early lexicality effects in adults and school-aged children (e.g., Kimppa et al., 2015; 2016) while the later time-window was chosen based on previous studies on word recognition in infants (e.g., Mills et al., 1993, 1997, Mills et al., 2005; Aygün & Rämä, 2022). However, instead of using peak-values for amplitude measures, we extracted amplitudes over two fixed time windows, which is a more typical procedure in infants’ studies. The analysis of variance (ANOVA) included four within-subject factors: *Word* (word, pseudoword), *Phase* (early, middle, late), *Recording Area* (frontal, parietal) and *Laterality* (left, right). The mean amplitudes extracted from **8** electrodes over each recording site were included in the statistical analyses: 12, 18, 19, 20, 23, 4 (F3), 27, and 28, in the left frontal, 3, 4, 5, 10, 117, 118, 123, and 124 (F4) in the right frontal, 52 (P3), 53, 59, 60, 61, 65, 66, and 67 in the left posterior 77, 78, 84, 85, 86, 90, 91, and 92 (P4) in the right posterior recording sites. The statistical analyses were conducted with SPSS (IBM SPP statistics, version 23) and the Greenhouse-Geisser correction was applied for non-sphericity when appropriate. Only main effects of Word and Phase and their interactions with other factors are reported here.

## Results

The ERPs in response to familiar words and novel pseudowords are illustrated over the two pairs of electrodes over the frontal and parietal recording sites (Fig. 1). The second syllable (e.i., disambiguation point) elicited a positive deflection (P100) around 100 ms after the disambiguation point followed by a negative-going deflection (N200-500). Amplitudes of the P100 in response to pseudowords decresed during the experiment while no modulation was observed for the N200-500. No modulations were detected in either neural response for familiar words. These findings were verified through statistical analyses.

**Fig. 1.**
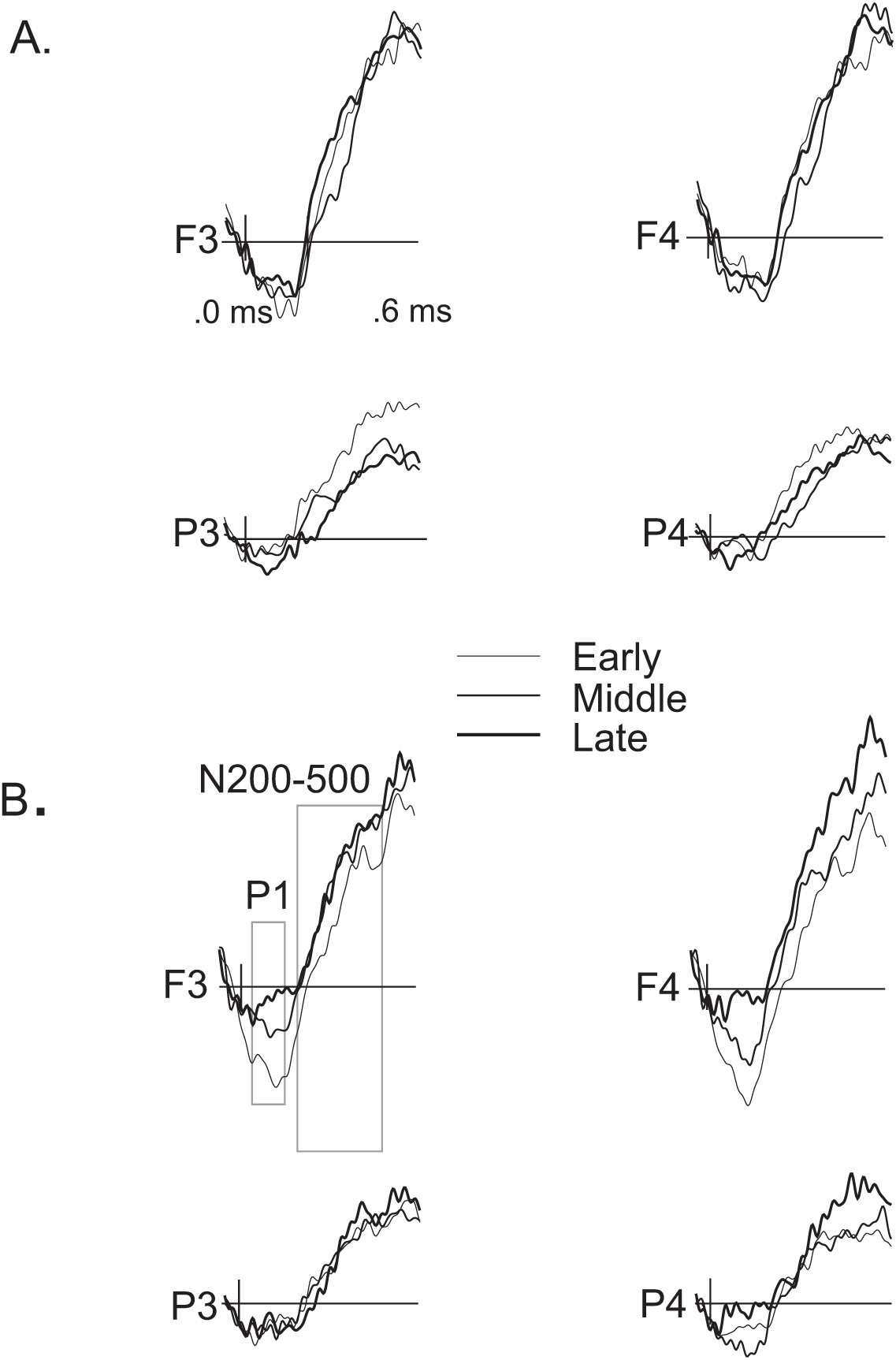
A. Grand-averaged waveforms for familiar words during the early, middle, and late phases of the experiment illustrated over two frontal (F3 and F4) and parietal (P3 and P4) electrodes. B. Grand-averaged waveforms for pseudowords words during the early, middle, and late phases of the experiment illustrated over two frontal (F3 and F4) and parietal (P3 and P4) electrodes. The vertical lines illustrate the word recognition point (e.i., second syllable). The gray rectangles indicate the time windows of the P100 and the N200-500 responses.

A significant main effect of Phase, F(2,48) = 3.52, p = .037, η² = .128 and a Word Type × Phase interaction, F(2,48) = 4.57, p = .016, η² = .160 were observed for P100 amplitudes. For pseudowords, P1 amplitudes were significantly reduced in the late compared to middle, (24) = 2.54, p = .018, Hedge’s g = 0.67 and early phases, t(24) = 3.27, p = .003, Hedge’s g = 0.79, whereas no difference was found between the early and middle phases, t(24) = 0.61, p = .549. In the late phase, P1 amplitudes were significantly smaller for pseudowords than for familiar words, t(24) = 2.83, p = .009, Hedge’s g = 0.51).

The N200–500 component was not significantly modulated by Phase, F(2,48) = 1.14, p = .325, η² = .046, Word Type, F(1,24) = 0.09, p = .772, η² = .004, or their interaction, F(2,48) = 1.51, p = .232, η² = .059.

**Fig. 2.**
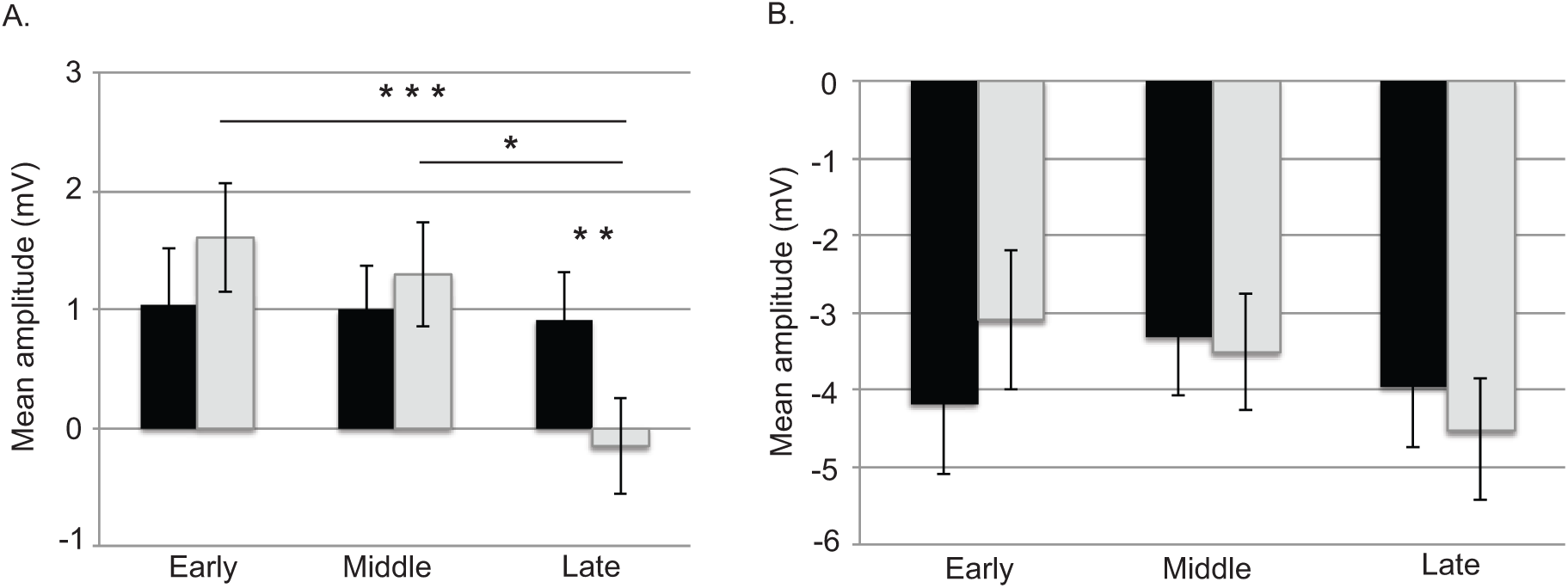
A. Mean amplitudes for P100 in response to familiar words (black bars) and pseudowords (grey bars) during the early, middle, and late phases of the experiment. B. Mean amplitudes for N200-500 in response to familiar (black bars) and pseudowords (grey bars) during the early, middle, and late phases of the experiment. Vertical bars indicate SEMs. *p < 0.05, **p < 0.01, ***p < 0.005

Changes in P1 amplitude for pseudowords across experimental phases were not significantly correlated with either receptive or productive vocabulary scores (all *r*s < .43, *p*s > .05). For the N200-500, however, there was a correlation between comprehensive vocabulary and amplitude change for pseudowords between the late and early phase, r = 0.553, p = 0.009. The lower the vocabulary was, the more negative the amplitudes were for the late phase compared with the early phase of the experiment.

## Discussion

This study investigated whether 24-month-old French-learning toddlers exhibit exposure-dependent neural changes to novel word forms without explicit training. Based on previous studies both in adults and school-aged children, we predicted changes in neural responses to pseudowords with exposure, modulated by vocabulary size. Our results demonstrated that the amplitudes of the P100, occurring around ∼100 ms after the word disambiguation point, decreased in response to pseudowords during the course of the experiment. Importantly, this reduction was not seen with familiar words. In contrast to our prediction, however, no correlation between amplitude changes and vocabulary skills was observed.

Earlier investigations show that brief passive exposure to meaningless pseudowords leads to an increase in early negativity with a latency of ∼50 ms both in the adult and school-aged children’s brain (e.g., Kimppa et al., 2015, 2016; Partanen et al., 2017; Leminen, Partanen, & Shtyrov, 2023) while in our study, we observed an early positive response, P100, which decreased selectively for pseudowords over the course of repeated exposure. The P100 component has been earlier associated with processing of acoustic features in speech (Čeponiene, Rinne, & Näätänen, 2002) and attentional processing of words (Mills et al., 2004). In infancy, both negative (e.g., Cheour, Ceponiene, Lehtokoski, Luuk, Allik, Alho, & Näätänen, 1998; Alho, Sainio, Sajaniemi, Reinikainen, & Näätänen, 1990) and positive (e.g., Leppänen, Eklund, & Lyytinen, 1997; Winkler, Kushnerenko, Horváth, Čeponienė, Fellman, Huotilainen, & Sussman, 2003) mismatch responses for deviant sounds have been found using oddball paradigms. These and some other recent findings on word recognition propose that polarity differences in general may reflect developmental changes in neural maturation: negative responses tend to emerge with age, becoming more characteristic of older infants and adults, while younger infants more commonly show positive responses (e.g., Männel & Friederici, 2013; von Holzen et al., 2018). Our study shows a modulation of positive response, P100, during the course of the experiment.

Despite earlier findings showing that early neural enhancement correlates with explicit learning outcomes in adults (Kimppa et al., 2015) and reading-related skills in children (Partanen et al., 2017), we did not find a correlation between P100 amplitude reduction in pseudowords and individual vocabulary skills, as measured by a parental vocabulary checklist. One interpretation of our findings is that unfamiliar novel word forms initially capture greater attentional resources, producing a larger early positivity, which then decreases as the forms become familiar through repeated exposure, possibly reflecting a form of neural habituation of the new word form. This interpretation is consistent with the finding that the amplitude reduction was specific to pseudowords: familiar words, already well enough represented in toddlers’ lexicons, did not show the same amplitude reduction. The difference in amplitudes for known words and pseudo words was significant only at the late phase of the task. This suggests that toddlers needed a certain amount of exposure before they showed a discriminative response between correct word forms and mispronounced forms.

The amplitude enhancement has been shown to occur faster in school-aged children (within approximately 4 minutes) than in adults (approximately 20–30 minutes) (Partanen et al., 2017), suggesting that the developing brain may be particularly sensitive to rapid reorganization in response to new phonological input, possibly reflecting heightened neural plasticity during periods of intensive language acquisition. In our study, we compared amplitudes across the initial, middle, and late phases of the experiment (each lasting approximately 4 minutes) and observed a significant reduction in early positivity between the initial and middle *versus* late phases, indicating that a notable change occurred within 8 to 12 minutes. This suggests that the signal change in 24-month-old toddlers was slightly slower than in school-aged children but still faster than in adults, suggesting that even though toddlers are at a period of intense lexical growth, the cortical mechanisms underlying rapid word form encoding are still maturing relative to those of older children.

In contrast to our expectation and previous findings (e.g., Mills et al., 1993, 1997; Aygün & Rämä, 2022), familiar words did not elicit more negative responses than meaningless pseudowords in the 200–500 ms time window. Furthermore, no significant amplitude changes, for either words or pseudowords, were observed in this later time window across the course of the experiment. The activity in the 200–500 ms window is typically associated with the processing of word meaning and familiarity (Mills et al., 1993, 1997), the learning of labels for novel objects (Mills et al., 2005; Torkildsen et al., 2008), and lexical-semantic processing more broadly (Rämä et al., 2013; Sirri & Rämä, 2015; Montgomery, Guellai, & Rämä, 2024). The lack of a familiarity effect in this time window may partly reflect the passive nature of the task. Our participants were not exposed to explicit links between words and objects, or meaningful links between word pairs, and thus, the N200–500 was not differentially elicited for familiar words and pseudowords. Interestingly, however, children with lower vocabulary skills showed more pronounced negative N200–500 amplitudes towards the end of the experiment for pseudowords, even though no meaning was assigned to the new word forms. One tentative interpretation is that toddlers with smaller vocabularies may allocate more sustained neural resources to processing unfamiliar phonological forms, perhaps because these forms are more difficult to categorize or integrate with existing lexical knowledge.

In conclusion, our results demonstrate that 24-month-old toddlers exhibit a P100 that diminishes with repeated exposure to novel, meaningless lexical items during a passive listening task. From a methodological standpoint, our results highlight the importance of monitoring within-session signal changes in infant ERP research: neural responses can be substantially modulated even within a short experimental session, and treating recordings as stationary across time may obscure meaningful exposure- or learning- related dynamics. Taken together, our findings suggest that the developing brain exhibits rapid, experience-dependent plasticity in response to novel input, even at early ERP latencies.

## Acknowledgements

We thank all families for their participation and contribution to this research. This work was supported by PME DIM Cerveau et Penśee, by the Institut Universitaire de France, and by a public grant overseen by the IdEx Université Paris Cité as part of the Labex Empirical Foundations of Linguistics - EFL [ANR-18-IDEX-0001].

## References

Alho, K., Sainio, K., Sajaniemi, N., Reinikainen, K., & Näätänen, R. (1990). Event-related brain potential of human newborns to pitch change of an acoustic stimulus. Electroencephalography and Clinical Neurophysiology/Evoked Potentials Section, 77(2), 151–155. 10.1016/0168-5597(90)90031-8

Akhtar, N., Jipson, J., & Callanan, M. A. (2001). Learning words through overhearing. Child development, 72(2), 416–430. 10.1111/1467-8624.00287

Aygün, O., & Rämä, P. (2022). Brain signatures of native and non-native words in French-learning 24-month-olds: The effect of vocabulary skills. Journal of Neurolinguistics, 62, 101055. 10.1016/j.jneuroling.2021.101055

Čeponiene, R., Rinne, T., & Näätänen, R. (2002). Maturation of cortical sound processing as indexed by event-related potentials. Clinical Neurophysiology, 113(6), 870–882. 10.1016/S1388-2457(02)00078-0

Cheour, M., Ceponiene, R., Lehtokoski, A., Luuk, A., Allik, J., Alho, K., & Näätänen, R. (1998). Development of language-specific phoneme representations in the infant brain. Nature neuroscience, 1(5), 351–353. 10.1038/1561

Fenson, L., Dale, P., Reznick, S., Thal, D., Bates, E., Hartung, J., Pethick, S., & Reilly, J. (1993). MacArthur Communicative Development Inventories: User’s guide and technical manual. Singular Publishing Group.

Gratton, G., Coles, M. G., & Donchin, E. (1983). A new method for off-line removal of ocular artifact. Electroencephalography and Clinical Neurophysiology, 55(4), 468–484. 10.1016/0013-4694(83)90135-9

Kimppa, L., Kujala, T., Leminen, A., Vainio, M., & Shtyrov, Y. (2015). Rapid and automatic speech-specific learning mechanism in human neocortex. NeuroImage, 118, 282–291. 10.1016/j.neuroimage.2015.05.098

Kimppa, L., Shtyrov, Y., Partanen, E., & Kujala, T. (2016). Read it again: Brain indices of learning novel words during reading. Scientific Reports, 6, 23492. 10.1038/srep23492

Leminen, A., Partanen, E., & Shtyrov, Y. (2023). Automatic word learning in adult brain: Cortical indices and time courses. Brain and Language, 236, 105220. 10.1016/j.bandl.2022.105220

Leppänen, P. H., Eklund, K. M., & Lyytinen, H. (1997). Event**-**related brain potentials to change in rapidly presented acoustic stimuli in newborns. Developmental Neuropsychology, 13(2), 175–204. 10.1080/87565649709540677

MacGregor, L. J., Pulvermüller, F., Van Casteren, M., & Shtyrov, Y. (2012). Ultra-rapid access to words in the brain. Nature Communications, 3, 711. 10.1038/ncomms1715

Männel, C. & Friederici, A. (2013). Accentuate or repeat? Brain signatures of developmental periods in infant word recognition. Cortex, 49, 2788–98. 10.1016/j.cortex.2013.09.003

Mills, D. L., Coffey-Corina, S. A., & Neville, H. J. (1993). Language acquisition and cerebral specialization in 20-month-old infants. Journal of Cognitive Neuroscience, 5(3), 317–334. 10.1162/jocn.1993.5.3.317

Mills, D. L., Coffey-Corina, S., & Neville, H. J. (1997). Language comprehension and cerebral specialization from 13 to 20 months. Developmental Neuropsychology, 13(3), 397–445. 10.1080/87565649709540685

Mills, D. L., Prat, C., Zangl, R., Stager, C. L., Neville, H. J., & Werker, J. F. (2004). Language experience and the organization of brain activity to phonetically similar words: ERP evidence from 14-and 20-month-olds. Journal of Cognitive Neuroscience, 16(8), 1452–1464. 10.1162/0898929042304697

Mills, D. L., Plunkett, K., Prat, C., & Schafer, G. (2005). Watching the infant brain learn words: Effects of vocabulary size and experience. Cognitive Development, 20(1), 19– 31. 10.1016/j.cogdev.2004.07.001

Montgomery, L., Guellai, B., & Rämä, P. (2024). Neural correlates of word learning in bilingual and monolingual 24-month-old infants. Brain and Language, 248, 105352. 10.1016/j.bandl.2023.105352

Partanen, E., Leminen, A., de Paoli, S., Bundgaard, A., Kingo, O. S., Krøjgaard, P., & Shtyrov, Y. (2017). Flexible, rapid and automatic neocortical word form acquisition mechanism in children as revealed by neuromagnetic brain response dynamics. NeuroImage, 155, 450–459. 10.1016/j.neuroimage.2017.03.066

Rämä, P., Sirri, L., & Serres, J. (2013). Development of lexical–semantic language system: N400 priming effect for spoken words in 18- and 24-month-old children. Brain and Language, 125(1), 1–10. 10.1016/j.bandl.2013.01.009

Shtyrov, Y., & MacGregor, L. J. (2016). Automatic ultrarapid activation and inhibition of cortical motor systems in spoken word comprehension. Proceedings of the National Academy of Sciences, 113(27), E3918–E3927. 10.1073/pnas.1524100113

Sirri, L., & Rämä, P. (2015). Cognitive and neural mechanisms underlying semantic priming during language acquisition. Journal of Neurolinguistics, 35, 1–12. 10.1016/j.jneuroling.2015.01.003

Torkildsen, J. V. K., Svangstu, J. M., Hansen, H. F., Smith, L., Simonsen, H. G., Moen, I., & Lindgren, M. (2008). Productive vocabulary size predicts event-related potential correlates of fast mapping in 20-month-olds. Journal of cognitive neuroscience, 20(7), 1266–1282. 10.1162/jocn.2008.20087

Von Holzen, K., Nishibayashi, L.-L., & Nazzi, T. (2018). Consonant and vowel processing in word form segmentation: An infant ERP study. Brain Sciences, 8: 24. 10.3390/brainsci8020024

Winkler, I., Kushnerenko, E., Horváth, J., Čeponienė, R., Fellman, V., Huotilainen, M., … & Sussman, E. (2003). Newborn infants can organize the auditory world. Proceedings of the National Academy of Sciences, 100(20), 11812–11815. 10.1073/pnas.2031891100

